# Alignment, calibration, and validation of an adaptive optics scanning laser ophthalmoscope for high-resolution human foveal imaging

**DOI:** 10.1101/2022.10.04.510799

**Authors:** Benjamin Moon, Martina Poletti, Austin Roorda, Pavan Tiruveedhula, Soh Hang Liu, Glory Linebach, Michele Rucci, Jannick P. Rolland

## Abstract

Advances in adaptive optics scanning laser ophthalmoscope (AOSLO) technology have enabled cones in the human fovea to be reliably resolved, providing new insight into human foveal anatomy, visual perception, and retinal degenerative diseases. These high-resolution ophthalmoscopes require careful alignment of each optical subsystem to ensure diffraction-limited imaging performance, which is necessary for resolving the smallest foveal cones. This paper presents a systematic and rigorous methodology for building, aligning, calibrating, and testing an AOSLO designed for imaging the cone mosaic of the central fovea in humans with cellular resolution. This methodology uses a two-stage alignment procedure and thorough system testing to achieve diffraction-limited performance. Results from retinal imaging of healthy human subjects show that the system can resolve cones at the very center of the fovea, the region where the cones are smallest and most densely packed.

## 1. Introduction

The use of adaptive optics in instrumentation for studying the human visual system has contributed to significant advances in understanding human retinal anatomy. Since the first use of adaptive optics in a retinal imaging system [1] and the subsequent application of adaptive optics to the scanning laser ophthalmoscope [2], the ability to compensate for the eye’s monochromatic aberrations has proved essential for achieving cellular resolution of retinal structures in the living human eye. Recent advances in the adaptive optics scanning laser ophthalmoscope (AOSLO) have enabled resolution of the smallest cones in the human fovea, providing a means to investigate the characteristics of the cone mosaic with unprecedented detail [3–8]. High-resolution AOSLO systems can also provide key data for the detection and monitoring of retinal degenerative diseases [9–11]. These promising applications of AOSLO depend on hardware that can reliably resolve the smallest human foveal cones, which requires an optical system that has simultaneously a sufficiently large numerical aperture at the retina to meet the resolution requirement and is well-corrected to meet the diffraction limit.

While AOSLO systems provide great potential for both basic and translational research, the complexity of these systems requires an optics expert for installation, alignment, and maintenance. A common AOSLO architecture for imaging the central fovea utilizes a series of four afocal telescopes to relay the entrance pupil of the system onto a fast (resonant) scanner, a slow (galvo) scanner, a deformable mirror, and the entrance pupil of the eye [3,12,13]. The afocal relay telescopes enable scanning and wavefront correction to occur at conjugate pupil planes, thus reducing scanning distortion artifacts and wavefront correction errors. Alignment errors in any of the telescopes can degrade the imaging performance by introducing optical aberrations. Imaging applications that resolve the smallest cones in the human central fovea require well-aligned optical systems because the diffraction-limited spot size for commonly used near-infrared imaging wavelengths is comparable to the smallest cone diameters.

Several publications have described the optical, mechanical, and system design of AOSLO systems [2,3,12–17]. Less emphasis has, however, been given to the alignment and system validation procedures used during the implementation of these AOSLOs. Alignment techniques for AOSLO systems utilizing a printed stencil and a shear plate to check collimation are implied in some publications [18–21], but rigorous documentation and systematic procedures are lacking in the literature. Yet these procedures are critically important for broadening access to the technology and enabling maintenance of the devices, especially in laboratories with limited optics expertise. Notable exceptions are provided by [22], which describes an active alignment strategy for minimizing the defocus after each relay telescope in the AOSLO, and by Lu *et al*. which presents an approach for characterizing the performance of the completed AOSLO by imaging a three-bar resolution target [23]. Nevertheless, a comprehensive approach to AOSLO implementation—including all aspects of assembly, alignment, and testing—is needed.

This paper presents rigorous methods for assembly, alignment, and system validation specifically aimed to enable greater access to high-resolution AOSLO technology. Active alignment strategies are used to ensure that each relay telescope is properly aligned in an afocal configuration, with collimated input light yielding collimated output light. A portable Shack-Hartmann wavefront sensor is positioned at each intermediate pupil plane during fine alignment of the system, allowing the wavefront to be measured after each relay telescope. The wavefront sensor is also positioned at the eye pupil plane and the system exit pupil planes to verify that both the light delivery and light collection paths are diffraction-limited according to the Maréchal criterion (i.e., RMS wavefront error less than 0.07 waves at any wavelength of operation). Further system validation tests include imaging a distortion grid target to measure and digitally correct for the sinusoidal distortion from the resonant scanner and imaging a three-bar resolution target to quantify the system resolution. Finally, human retinal images are collected, demonstrating that the smallest cones at the center of the fovea are resolved.

## 2. Implementation and alignment

### 2.1 System design and specifications

The design for the AOSLO used in this study is optimized for diffraction-limited performance over a 1-degree square field of view. The design closely resembles the system described in Mozaffari *et al*. [13]. The schematic for the system is shown in Fig. 1 and the specifications are listed in Table 1.

**Fig. 1:**
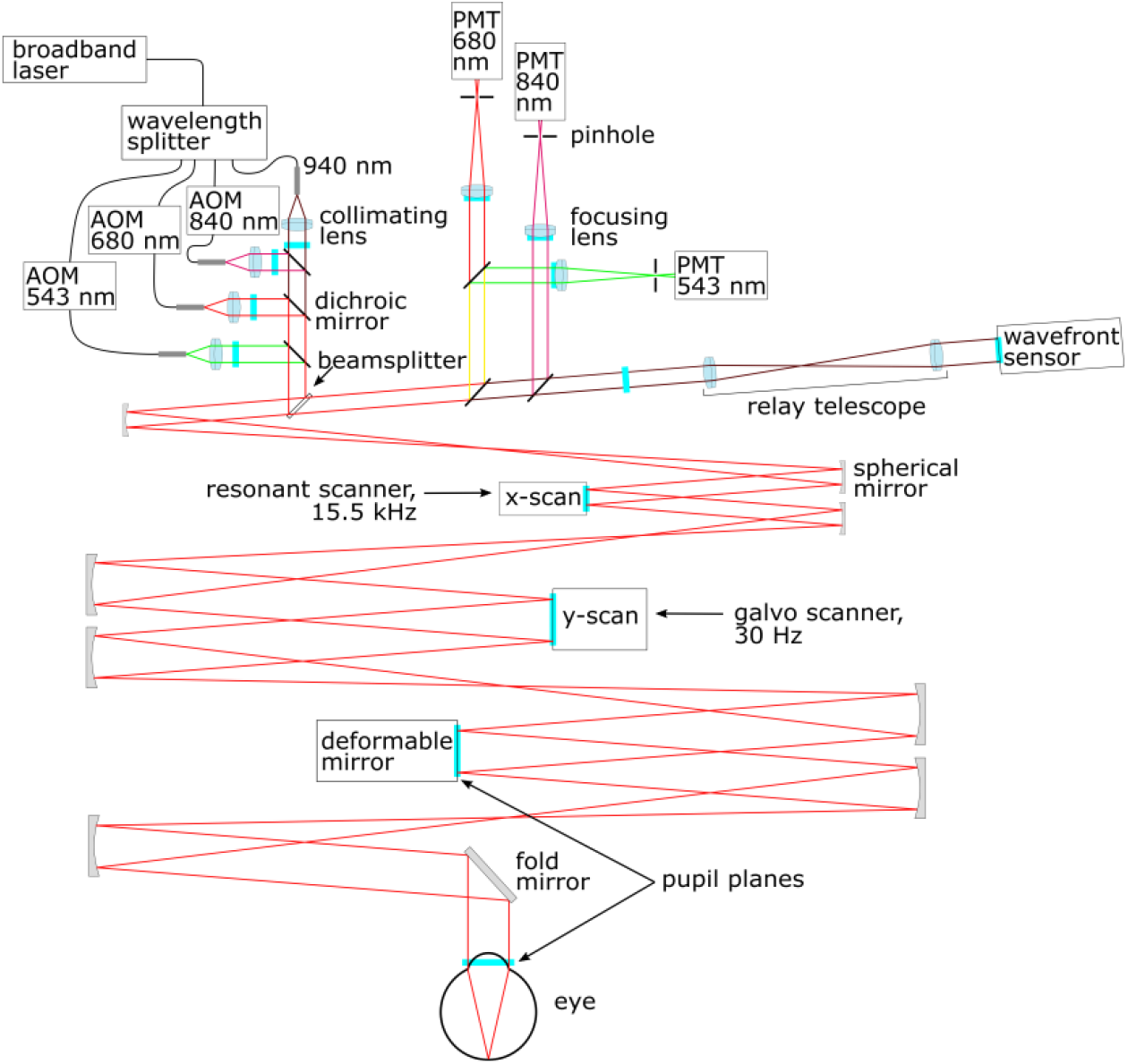
Schematic for the AOSLO system showing the light delivery path, the scanning system, the detection arm, and the wavefront sensing arm. The pupil planes for the system are shown as cyan lines. The mirrors for each telescope are spherical, and they are arranged in a non-planar configuration to minimize astigmatism.

**Fig. 2:**
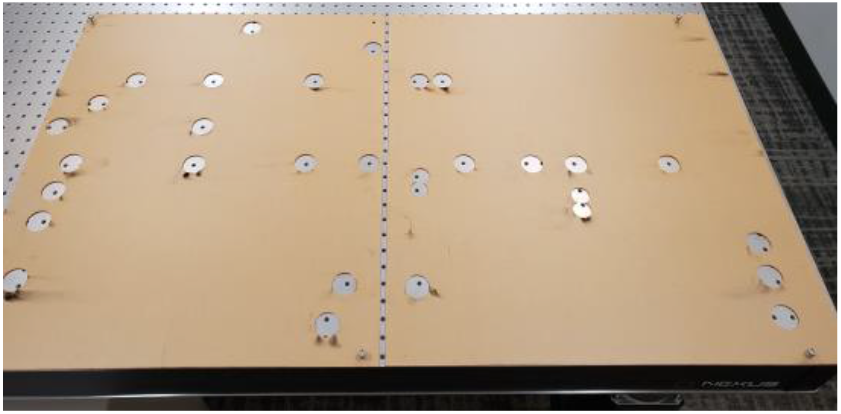
Laser-cut stencil used to transfer the post coordinates onto the optical table during the assembly and coarse alignment of the AOSLO.

**Table 1:** System specifications for the AOSLO.

| System component | Specifications | Notes |
| --- | --- | --- |
| Light source | Supercontinuum fiber laser, 470-2000 nm, 6 W output power | SuperK Extreme, EXR-20 (NKT Photonics A/S, Birkerød, Denmark) |
| Spectral filtering | $543 \pm 11$, $680 \pm 11$, $840 \pm 6$, $940 \pm 5$ nm (HWHM) | Fiber-coupled wavelength splitting device with Thorlabs optomechanics and bandpass filters from Semrock |
| Acousto-optic modulators (AOMs) | 0-50 MHz modulation, fiber-coupled AOMs | TEM-210-50-10-543/-680/-840-2FP-SM (Brimrose Corp., Sparks Glencoe, MD) |
| Horizontal scanner | 15.5 kHz resonant scanner, Ø4.5mm active area | SC-30 (Electro-Optical Products Corp., Queens, NY) with custom mirror from Optimax |
| Vertical scanner | Galvanometer scanner, Ø9.5mm active area | 6SD11170 (Cambridge Technology, Bedford, MA) |
| Photomultiplier tubes (PMTs) | Ø5 mm sensing area, 1 ns rise time, GaAs/GaAsP | H7422-40/-50 (Hamamatsu, Shizuoka Pref., Japan) |
| Amplifiers for PMTs | 0-50 MHz transimpedance amplifier, 25 mV/μA gain | C6438-01 (Hamamatsu, Shizuoka Pref., Japan) |
| Confocal pinholes | Ø40/50 μm precision pinholes placed in front of PMTs | P40D/P50D (Thorlabs, Newton, New Jersey) |
| Wavefront sensor (WFS) camera | 1.4 MP (1384 x 1032 pixels), 45 fps, 2/3" CCD | GS3-U3-15S5M-C (FLIR, Wilsonville, OR) |
| WFS lenslet array | 200 μm pitch, 9 mm focal length, 12 x 12 mm size | APO-Q-P200-R4.15 (Advanced Microoptic Systems GmbH, Saarbrücken, Germany) |
| Deformable mirror | 97 actuators, Ø7.2 mm pupil, 1.5 ms settling time | DM97-08 (ALPAO, Montbonnot, France) |
| Pupil diameter (at eye) | 7.2 mm | Size set by deformable mirror aperture |
| Adaptive optics closed-loop frequency | 15-25 Hz | - |
| Imaging frame rate | 30.3 fps | Set by the horizontal scanner frequency |
| Imaging field of view (FOV) | 1 degree by 1 degree square | Adjustable up to 1.5 degrees with 1 degree being typical. Measured at the eye pupil plane |
| Imaging resolution | 512 x 512 pixels, 8.53 pixels/arcmin @ 1° FOV | - |

The system has four spectral channels with central wavelengths of 543 nm, 680 nm, 840 nm, and 940 nm. Three of the spectral channels—543 nm, 680 nm, and 840 nm—are configured for retinal imaging and stimulus delivery, enabling flexible experiment design. Each of these three channels is equipped with a PMT for recording the backscattered light from the retina and accurately localizing the stimulus position on the retina, which is essential for studying fixational eye movements and visual perception in AOSLO psychophysics experiments. The 940 nm channel is used for wavefront sensing. The use of this independent wavefront sensing channel at 940 nm enables all the backscattered light from the imaging channels to go to the imaging detection path instead of being split between imaging and wavefront sensing. At 940 nm, this light is invisible to the human eye, and the retinal exposure limits are less stringent compared with the other imaging and stimulus delivery channels.

These four spectral channels are produced by dividing and spectrally filtering the broadband output from a supercontinuum fiber laser. The three imaging and stimulus delivery channels (i.e., 840 nm, 680 nm, and 543 nm) are modulated with acousto-optic modulators (AOMs), allowing the intensity of each channel to be independently adjusted and enabling stimuli to be drawn on the retina by embedding visible light patterns in the AOSLO raster scan. The wavefront sensing channel (i.e., 940 nm) is not modulated, which results in a wavefront correction that is averaged over the field of view as the beacon is scanned across the retina. This field-averaged correction is preferable for the chosen application where a small field of view (e.g., 1-degree square) is used, which is smaller than the isoplanatic patch size of the human eye [24,25]. For other applications using a larger field of view, a stationary beacon for on-axis wavefront correction may be a better choice [16,17]. Fast scanning is achieved by a 15.5 kHz resonant scanner. Slow scanning uses a 30 Hz galvo scanner. The scanning field of view can be adjusted by changing the control voltages for each scanner. The optical design uses spherical mirrors in an off-axis and out-of-plane configuration to achieve an unobscured system with minimal astigmatism [3].

The adaptive optics (AO) control loop consists of a custom Shack-Hartmann wavefront sensor, a deformable mirror with 97 actuators, and custom software for wavefront control. When the adaptive optics control loop is running, the aberrations of the human eye are corrected, resulting in a diffraction-limited spot of light being scanned across the retina. Backscattered light then travels back through the scanning system and is separated into different spectral channels using dichroic filters in the detection arm. The light in each of the imaging channels is focused onto a sub-Airy-disk confocal pinhole by a focusing lens and then detected by a high-sensitivity photomultiplier tube (PMT).

### 2.2 Coarse alignment strategy

A two-stage assembly and alignment procedure was used to efficiently install and align the components of the AOSLO while targeting the demanding imaging performance requirements. Section 2.2 describes the coarse alignment while Section 2.3 describes the fine alignment. Following the technique described in [13], a stencil containing all of the post locations on the optical table was generated from the optomechanical CAD model for the AOSLO. This stencil was laser cut in a thin acrylic sheet in two pieces and was then used to transfer the component post locations onto the optical table with a position uncertainty of less than +/- 0.5 mm. This approach simplified the coarse alignment procedure and enabled the system to be rapidly assembled on the optical table. The component heights were determined from the CAD model of the system and were realized by measuring the distance between the surface of the optical table and the corresponding reference plane on the mechanical mounts. Using this approach, each optical component was aligned in three dimensions with a position uncertainty of +/- 1 mm, which includes tolerances on the mechanical mounting holes and hardware.

After aligning the position of each component on the optical table, the angle of each mirror in the scanning system was optimized to center the beam on the next mirror in the system. Precision-machined alignment guides (LMR1AP and LMR2AP from Thorlabs) were used to visualize the center of each spherical mirror (see Fig. 3). The 543 nm channel was used as the primary alignment channel because the green light could be easily seen at low power levels (i.e., less than 1 mW). Using the alignment guide, the beam was centered on the distal mirror in the beam path with an uncertainty of +/- 0.5 mm. Centering the beam on the distal mirror is important as it will set the angular precision of the prior mirror along the path. The +/- 0.5 mm beam-centering uncertainty on the distal mirror corresponds to a maximum angular misalignment of +/- 7 arcmin for the prior mirror with the shortest optical path length to the distal mirror. From conducting a tolerancing and sensitivity analysis of the AOSLO using optical design software, it was determined that the system would maintain diffraction-limited performance for misalignments of up to +/- 30 arcmin for this mirror. Therefore, the angular alignment attainable with the alignment guides is well within the alignment tolerance.

**Fig. 3:**
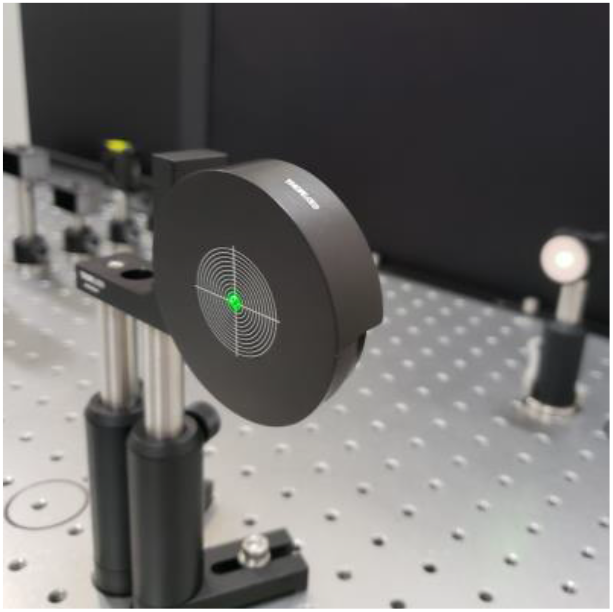
Alignment guide used to center the laser beam on each of the spherical mirrors in the scanning system. Concentric circles surround a central hole, with a spacing of 1 mm between consecutive lines. Using these alignment guides, angular alignment better than +/- 7 arcmin can be achieved for each of the mirrors in the system. The laser beam from the 543 nm channel is shown being used to align a pair of mirrors in the scanning system.

### 2.3 Fine alignment strategy

After completing the coarse alignment, a portable Shack-Hartmann wavefront sensor (WFS40-7AR from Thorlabs) was sequentially placed at each intermediate pupil plane in the system during the fine alignment phase. This technique, as described by Steven [22], allows for the collimation to be checked after each afocal relay telescope. Alternative approaches have utilized a shear plate to check collimation [18–21]. The decision to use a Shack-Hartmann wavefront sensor instead of a shear plate was motivated by the temporal coherence properties of the light sources used in this multi-wavelength design. The bandwidths of the four spectral channels range from 10 nm to 22 nm FWHM, with coherence lengths between 4 μm and 27 μm. These short coherence lengths preclude the use of shear plates for checking collimation, which have optical path differences of several millimeters to tens of millimeters. Although an auxiliary alignment laser with a narrow linewidth could have been installed for use with a shear plate, using a wavefront sensor allows all channels to be tested with their as-used spectral content and removes a source of alignment error that would be introduced when switching between the alignment laser and the laser used for imaging and stimulus delivery.

This fine alignment process required that the scanners and deformable mirror be temporarily removed to enable proper positioning of the wavefront sensor at the pupil planes. After using the wavefront sensor to verify that the incident wavefront was collimated at the entrance pupil plane for each of the four wavelength channels, the wavefront sensor was placed at the first intermediate pupil plane, corresponding to the fast scanner location. The wavefront sensor was aligned to center the incident beam on the lenslet array and minimize tilt. Next, a wavefront measurement was collected, and the corresponding defocus was recorded. This initial measurement showed that there was 0.017 Diopters of defocus after the first telescope. As shown in Fig. 4, this measured defocus can be used to determine the required axial shift of mirror 2 to remove the residual defocus, as described by Eqs. (1) and (2). The axial position of mirror 2 was adjusted while keeping mirror 1 stationary to ensure that the entrance pupil of the telescope did not change during alignment.

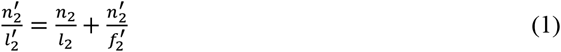

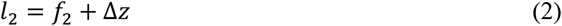

**Fig. 4:**
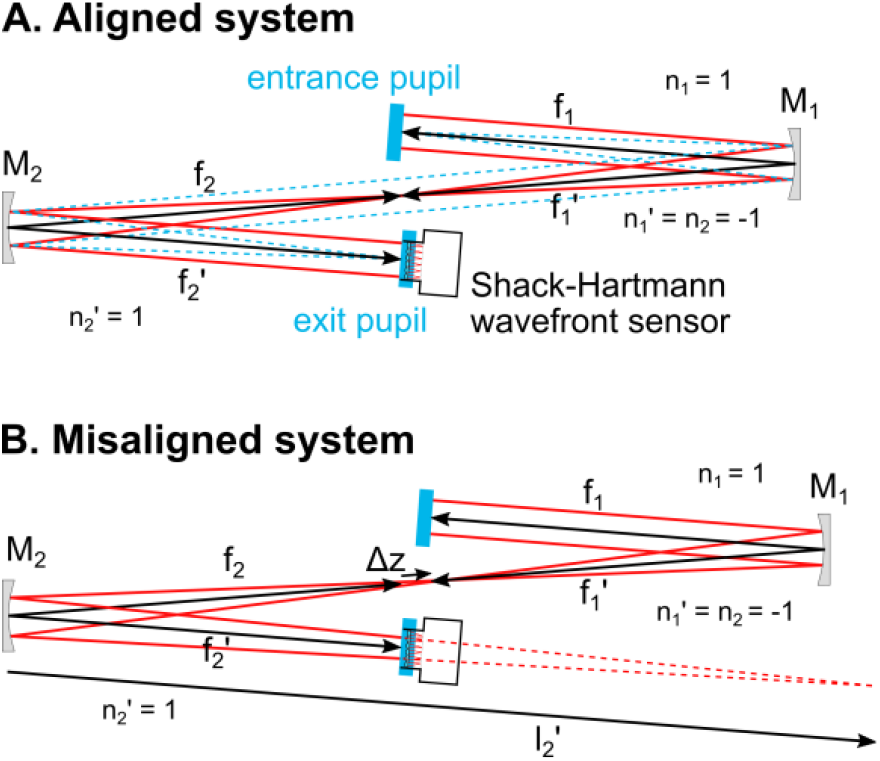
Simplified schematic for a single telescope demonstrating the wavefront measurement and adjustment procedure. For the aligned system, the wavefront sensor at the exit pupil plane measures a collimated wavefront. For the misaligned system—where there is an axial positioning error for mirror 2—the wavefront at the exit pupil plane is converging or diverging. By measuring the wavefront curvature at the exit pupil plane, the axial shift required to correct for the measured defocus can be determined. As the position of mirror 2 is adjusted, the wavefront at the exit pupil plane is continuously measured until the residual defocus is reduced below the alignment tolerance.

Eq. (1) describes the thin-lens imaging properties of mirror 2, where *n*_2_ and 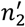 are the refractive indices before and after mirror 2 respectively, *l*_2_ is the object distance, 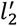 is the image distance, and 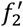 is the focal length of mirror 2. Eq. (2) shows that the object distance *l*_2_ can be replaced by the sum of the front focal length, *f*_2_, and the axial positioning error, Δ*z*. The object distance, 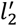, is determined from the wavefront measurement and is inversely proportional to the measured vergence, 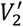, as shown in Eq. (3).

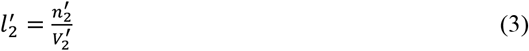

Using Eqs. (1), (2), and (3), the axial positioning error, Δ*z*, can be solved for. Recalling that 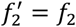 for a mirror, the resulting equation can be simplified, as shown in Eq. (4).

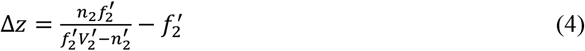

Plugging in the values corresponding to mirror 2 in the first telescope 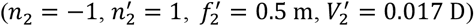, it is determined that the measured defocus corresponds to an axial positioning error of 4.3 mm for mirror 2. After adjusting the position of mirror 2 by 4 +/- 1 mm (moving mirror 2 closer to mirror 1), the measured defocus at the exit pupil of the relay telescope was 0.008 Diopters, which is within the alignment tolerance of +/- 0.01 D. This alignment tolerance of 0.01 D is a conservative measure of the sensitivity limit of the Shack-Hartmann wavefront sensor used to check the collimation of each telescope. The resulting wavefront error from this residual 0.01 D of defocus is less than λ/8 peak-to-valley for all wavelengths used in the system, which is well below the diffraction-limited criterion of λ/4 peak-to-valley for defocus. After optimizing the spacing between mirrors 1 and 2, the fast scanner was reinstalled and its position along the optical axis was optimized by measuring along the optical axis from mirror 2 to the fast scanner and verifying that the distance was equal to the focal length of mirror 2. With the first telescope realigned and the fast scanner reinstalled, the beam angles were once again optimized before proceeding to the next telescope.

The procedures described above for the first telescope were repeated in each of the other three telescopes to ensure optimum alignment. First, the wavefront sensor was placed at the pupil plane after a given telescope and a wavefront measurement was taken. Then the required mirror shift along the optical axis was calculated and implemented. After making the position adjustment to the mirror and the wavefront sensor to ensure that the wavefront sensor remained at the pupil conjugate, another measurement was taken to verify that the mirror shift properly compensated for the residual defocus in the telescope. Mirror angles were then optimized for beam centration throughout the whole scanning system. Once all four telescopes were aligned using this procedure, wavefront measurements were taken at the eye pupil plane. It is important to note that the deformable mirror was replaced with a flat mirror (PF10-03-P01 from Thorlabs) during this fine alignment procedure to ensure that the resting shape of the deformable mirror would not introduce unknown defocus or aberrations into the system.

This active alignment technique—where a portable Shack-Hartmann wavefront sensor measures the wavefront shape and corresponding defocus at a pupil plane while adjustments are made to the axial position of a lens or mirror—was also utilized to install the relay telescope in front of the wavefront sensor, as pictured in Fig. 1. First, a flat mirror was placed at the eye pupil plane to reflect the incident light back through the scanners toward the detection channels. The portable wavefront sensor was then placed at the exit pupil plane to verify that there was good agreement between the wavefronts measured at the eye pupil plane and the exit pupil plane for the wavefront sensing channel. After verifying that the wavefront at the exit pupil was flat (less than +/- 0.01 D of defocus) and diffraction-limited (RMS wavefront error less than 0.07 waves), the two lenses that make up the relay telescope were installed. Measurements with a ruler were used to ensure that the first lens was positioned one focal length away from the pupil plane and the two lenses were separated by the sum of their focal lengths corresponding to an afocal relay telescope. Next, the portable wavefront sensor was placed at the exit pupil plane of the relay telescope. There was some residual defocus in the wavefront measurement at this plane, which was removed by adjusting the axial position of the second lens while monitoring the wavefront measured at the exit pupil plane. Using this approach, non-common-path aberrations between the light delivery, imaging, and wavefront sensing paths were negligible and focusing was optimized, as demonstrated by the diffracted-limited wavefront measurements collected at the exit pupil planes.

## 3. Validation and Calibration

### 3.1 Wavefront measurements at the eye pupil plane

By using the fine alignment strategy described in Section 2.3, each telescope in the AOSLO was aligned to have less than +/- 0.01 Diopters of residual defocus. The portable Shack-Hartmann wavefront sensor was then placed at the eye pupil plane. Wavefront measurements at the eye pupil plane were collected for each of the four wavelengths used in the system. The deformable mirror was removed from the system during these tests and replaced with a flat mirror to ensure that the resting shape of the deformable mirror would not contribute toward the measured wavefront, just as was done during fine alignment. The beam diameter at the eye pupil plane is 7.2 mm, so all measurements were collected with a pupil diameter of 7.2 mm defined in the wavefront sensor software and a manually adjusted pupil offset to keep the measurement pupil centered on the beam. Raw wavefront maps were saved using the wavefront sensor software and were later analyzed in MATLAB (Natick, Massachusetts).

Fig. 5 shows the wavefront maps for each spectral channel at the on-axis field point. The RMS wavefront error is less than 0.05 waves for all channels, which is well below the Maréchal criterion for a diffraction-limited optical system (i.e., less than 0.07 waves). Results from modeling the AOSLO in optical design software show that the RMS wavefront error changes by less than 0.015 waves across the full field of view, so the on-axis measurement was determined to be representative of the performance across the field of view. There is a small amount of residual astigmatism, coma, and higher-order aberrations present in the wavefront maps, but these residual aberrations have a negligible impact on the image quality because the resolution and image quality are primarily limited by diffraction. The average peak-to-valley wavefront error is 0.27 waves across the four channels.

**Fig. 5:**
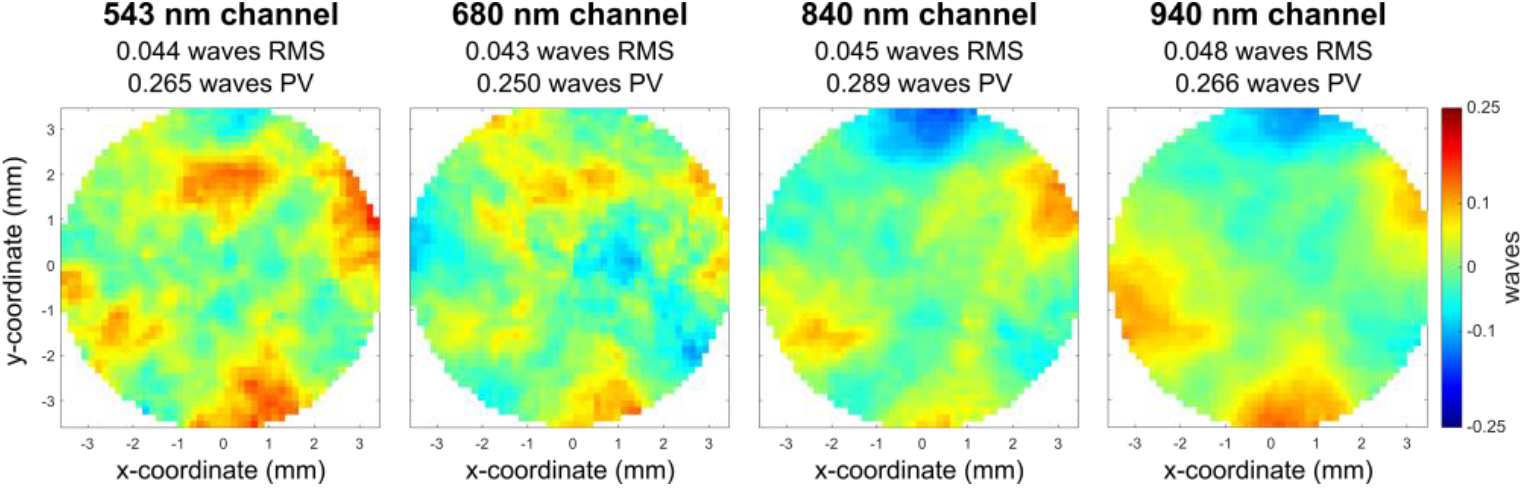
Wavefront measurements at the eye pupil plane for each of the four spectral channels. These results demonstrate that the system is well aligned and diffraction-limited according to the Maréchal criterion (i.e., RMS wavefront error less than 0.07 waves) for the on-axis field.

These measurements at the eye pupil plane validate the system alignment by showing that the combination of the individually aligned relay telescopes results in a diffraction-limited optical system. During the fine alignment of each relay telescope the defocus was minimized, but other aberrations—such as astigmatism and coma—were not addressed. Combining the relay telescopes together in an out-of-plane configuration minimizes the overall system astigmatism [3], resulting in a diffraction-limited optical system. Resolving the smallest foveal cones in the human retina requires this level of aberration correction. For the primary imaging wavelength of 680 nm and a standard 60-Diopter eye with a pupil diameter of 7.2 mm (i.e., the maximum pupil size attainable with this AOSLO), the Rayleigh resolution criterion for point-like objects is 1.9 μm (for 840 nm, it is 2.4 μm). This means that objects separated by 1.9 μm on the human retina will be just resolved when the system is limited by diffraction. Since the minimum center-to-center spacing for human cone photoreceptors is around 2.43 +/- 0.24 μm at the foveal center [26], the ability to resolve these smallest cones requires the optical system to be diffraction-limited for each spectral channel. By ensuring that the aligned optical system meets this requirement, the full range of adaptive optics correction available with the deformable mirror can be applied to correct for the aberrations of the human eye.

At this stage, the deformable mirror was installed to assess the functionality of the AO control loop. The setting for the best-flat shape of the deformable mirror was applied, achieving better than 0.03 waves RMS wavefront error. Positioning the portable wavefront sensor at the eye pupil plane was also beneficial for fine-tuning the longitudinal chromatic aberration (LCA) that was introduced into the system to pre-compensate for the average LCA of the human eye. Inherent uncorrected LCA of the human eye would cause the four different wavelengths to focus at different retinal depths if all wavelengths were collimated at the eye pupil plane [27–29]. To account for the 1.2 Diopters of LCA between the shortest and longest wavelengths used in the system (i.e., 543 nm and 940 nm, respectively), the technique described in [13] was employed, where the distance between the input fiber tip and collimating lens was adjusted to introduce vergence for different spectral channels. With the 680 nm imaging channel being collimated (i.e., 0 Diopters of vergence), the input fiber for the 543 nm channel was moved 3.9 mm closer to the collimating lens, which has a focal length of 40 mm. The fiber for the 840 nm channel was moved away from its collimating lens by 2.5 mm, and the fiber for the 940 nm channel was moved 3.5 mm away from its collimating lens. By continuously monitoring the wavefront curvature at the eye pupil plane during these system adjustments, the correct amount of LCA (i.e., 1.2 Diopters between 543 nm and 940 nm) was added to ensure that all wavelengths would focus at the same retinal plane for the average human eye. The wavefront sensor was also used to fine-tune the beam coalignment of the four spectral channels by measuring the wavefront tip and tilt at the eye pupil plane after making the LCA adjustments. The performance was diffraction-limited after making the LCA adjustments based on imaging a resolution target as further detailed in Section 3.3.

### 3.2 Distortion grid measurements and calibration

Measurement and calibration of the field of view was conducted using a distortion grid target (R1L3S3P from Thorlabs). A model eye lens, consisting of an achromatic doublet (AC254-050-B from Thorlabs), was placed with the front surface of the lens at the exit pupil plane (i.e., eye pupil plane) of the system. A paper target was placed at the back focal plane of the model eye lens, and the adaptive optics control loop was enabled to compensate for any residual system aberrations. Once stable correction was achieved, the adaptive optics control loop was switched off and the wavefront was monitored to ensure good correction was maintained. The paper target was then replaced with the distortion grid target, which was carefully aligned at the back focal plane of the lens to achieve high-contrast imaging. By following these procedures, adverse effects from the specular reflection of the distortion grid target were avoided. Imaging was conducted with the 840 nm channel. The scanner drive signal amplitudes were adjusted to control the field of view by setting a 10-bit digital number on a custom graphical user interface that communicated with the scanner drivers. Using the known grid spacing of the distortion grid (i.e., 50 μm grid spacing with +/- 1 μm spacing tolerance) and the focal length of the model eye lens, the full field of view in both the horizontal and vertical directions was determined from measurements of distortion grid images. Modeling in optical design software was conducted to verify that the model eye lens did not introduce significant distortion: the maximum distortion over a 1-degree square field of view was less than 20% of the pixel width. Eq. (5) was used to compute the full field of view in either the horizontal or vertical direction: *θ*_*FFOV*_ is the horizonal (respectively vertical) full field of view, *x* (respectively *y*) is the measured width (or height) of the distortion grid image, and *f*^*′*^ is the focal length of the model eye lens.

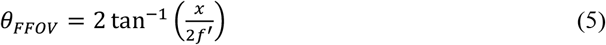

Calibration curves for both the horizontal and vertical scanners were generated by adjusting the scanner amplitudes to achieve seven different square fields of view ranging from 0.13 degrees to 1.5 degrees for the full field of view. Distortion grid images for each of the seven measurements are shown in Fig. 6A. For the horizontal scanner, a cubic polynomial fit was used, while a quadratic polynomial fit was used for the vertical scanner. Different polynomial orders were used for the two scanners to accurately capture the different amplitude responses of the two types of scanners. The measured data points and the polynomial fits are displayed in Fig. 6A, with red denoting the horizontal scanner and blue denoting the vertical scanner. The equations for the two polynomial fits are given by Eqs. (6) and (7), where *H* is the digital number for the horizontal scanner, *θ*_*H*_ is the horizontal full field of view in degrees, *V* is the digital number for the vertical scanner, and *θ*_*V*_ is the vertical full field of view in degrees.

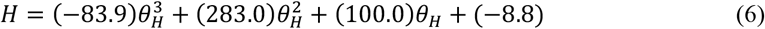

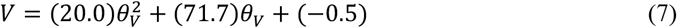

**Fig. 6:**
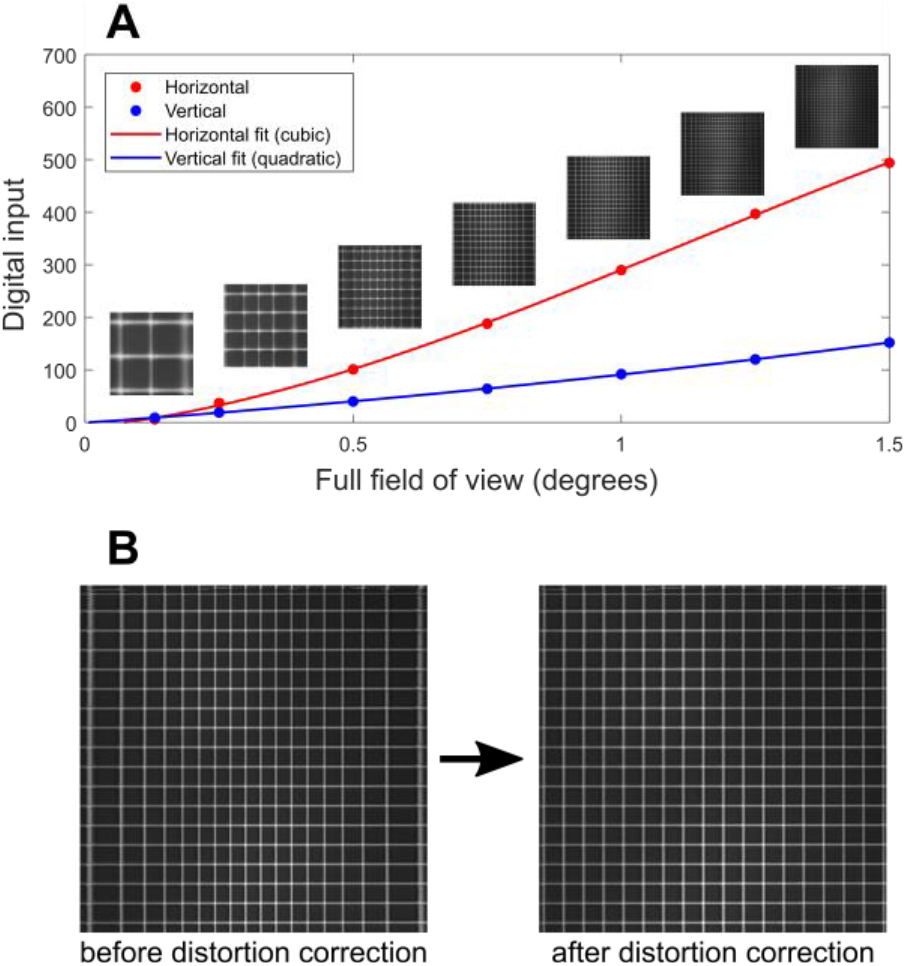
A distortion grid was used to calibrate the field of view and correct for the sinusoidal distortion from the resonant scanner. A: images were taken at seven different square field of view settings and used to generate calibration curves for both the horizontal and vertical scanners. B: the distortion grid image on the left shows stretching at the edges of the frame due to the sinusoidal distortion from the resonant scanner. After distortion correction (right), the grid image shows uniform spacing across the field of view.

These calibration curves provide a means for determining the correct amplitude settings for the two scanners to achieve a desired field of view. For a 1-degree square full field of view the required amplitude settings are *H* = 290 and *V* = 91.

After calibrating the field of view and setting the scanner amplitudes to achieve a 1-degree square full field of view, a lookup table was generated to remove the sinusoidal distortion in the image caused by the horizontal scanner. The horizontal scanner is a resonant scanner, which oscillates at a frequency of 15.5 kHz. Its motion characteristics are sinusoidal, which cause the image to be stretched at the left and right edges of the frame and compressed near the center. The left image in Fig. 6B shows the distortion grid image before correcting for the sinusoidal distortion. Stretching at the right and left edges of the image can be observed. By quantifying the location of each grid line in the image and comparing the real location to the expected location, the sinusoidal distortion can be corrected. A lookup table is then generated and used to resample the image pixels, which results in the distortion-free image in the right panel of Fig. 6B. This lookup table is then applied for real-time distortion correction in both the live and recorded videos. The distortion correction procedure is implemented in the data acquisition software, which has been described in prior art [14,30–32]. Using a grid target for distortion correction instead of a Ronchi Ruling is useful for correcting residual non-orthogonality of the scanners and enables simultaneous measurement of the horizontal and vertical field of view.

### 3.3 Resolution target measurements

After calibrating the field of view and removing the sinusoidal distortion, a 1951 USAF three-bar resolution test target (R1DS1P from Thorlabs) was imaged to quantify the resolution limit of the completed AOSLO. The resolution target was placed at the back focal plane of the model eye lens and was imaged with each of the three imaging channels. The resolution limits in both the horizontal and vertical directions were quantified as the maximum spatial frequency that could be resolved with all three bars visible in the image. The measured resolution limits were then compared to the Rayleigh resolution limit for the imaging setup to demonstrate that the system could resolve features up to the diffraction limit.

Imaging was conducted over a 1-degree square field of view over a period of 10 seconds following the same procedures outlined in Section 3.2: the AO loop was initialized with a paper target and then switched off. The 300 captured frames were then averaged to obtain the image in Fig. 7A. The expanded view in Fig. 7B shows that features are resolved up to group 7, element 2 for the horizontal axis and group 7, element 3 for the vertical axis. The Rayleigh resolution limit for this imaging configuration (i.e., wavelength of 840 nm, pupil diameter of 7.2 mm, and focal length of 50 mm) is 7.1 μm. This minimum feature separation for resolution corresponds to a maximum spatial frequency of 141 lp/mm. The spatial frequencies for elements 2 and 3 of group 7 are 144 and 161 lp/mm, respectively, which demonstrates that the imaging system can resolve features up to—and slightly beyond—the Rayleigh resolution limit.

**Fig. 7:**
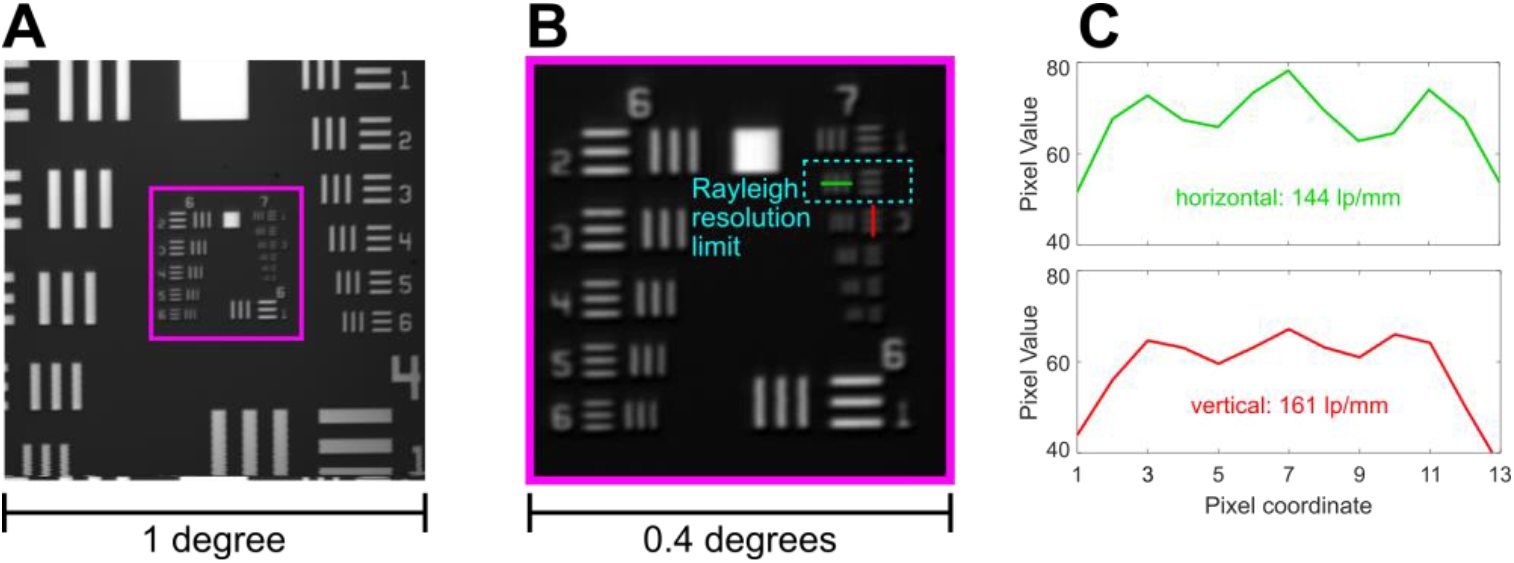
A resolution target was placed at the back focal plane of a model eye lens and imaged with the 840 nm channel. The results demonstrate that the system can resolve features up to the Rayleigh resolution limit. Imaging was conducted over a 1-degree square field of view, as shown in A. A close-up view of the central 0.4-degrees square is shown in B. Pixel value lineouts are shown in C for the maximum spatial frequency that is resolved for both the horizontal and vertical directions. The Michelson contrast is greater than 4% for each of the peaks shown.

The small discrepancy between the resolution in the horizontal and vertical directions is a consequence of the scanning architecture. Residual jitter in the hsync clock signal due to limitations of the custom data acquisition electronics can cause consecutive horizontal lines in the image to shift by +/- 1 pixel relative to the previous lines. When averaging across many frames, this horizontal jitter can reduce the contrast of the vertical bars used to quantify resolution along the horizontal axis. The horizontal jitter does not impact the contrast of the horizontal bars used to quantify vertical resolution because the direction of the jitter is along the length of the bars. This reasoning explains why the resolution for the red (vertical) line in Fig. 7B,C is higher than the resolution for the green (horizontal) line.

After imaging with the smallest features at the center of the field of view, the resolution target was shifted laterally to center the smallest features at each of the four corners of the field of view. The measured resolution at these locations was identical to the resolution measured at the center, as expected from the modeling results using optical design software. Resolution measurements were also conducted with the two visible channels, and the results are summarized in Table 2. The 680 nm channel exceeds the Rayleigh resolution limit. The Rayleigh resolution limit for the 543 nm channel is in between two of the features on the resolution target: element 5 of group 7 has a spatial frequency of 203 lp/mm and element 6 has a spatial frequency of 228 lp/mm, while the Rayleigh limit is 217 lp/mm. Element 5 is resolved, while element 6 is not resolved. Quantifying the baseline resolution of the system by imaging a resolution target validates the system performance and enables routine alignment and image quality assessments. The resolution limit is measured before each human imaging session to verify the system performance for whichever spectral channel is being used.

**Table 2:**
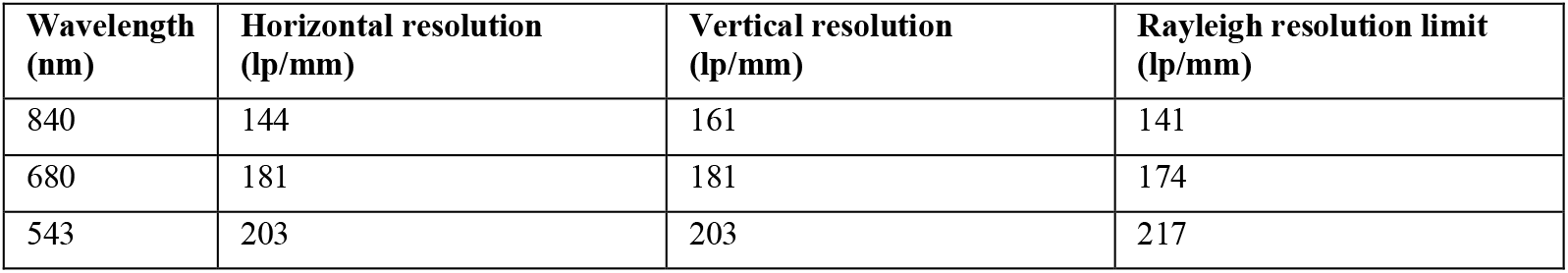
Resolution measurements for each of the three imaging channels.

## 4. Human retinal imaging

### 4.1 Imaging methods

To demonstrate the effectiveness of the alignment procedures described in this article, retinal imaging of seven subjects was conducted. Two subjects were emmetropic, two were hyperopic (+0.5 D), and three were myopic (−1 D to -3.5 D range). The imaging protocols adhered to the ethical standards of the Research Subjects Review Board (RSRB) at the University of Rochester, and this study was approved by the RSRB. All subjects provided informed consent to participate in the study. One drop each of 1% tropicamide and 2.5% phenylephrine ophthalmic solutions were administered to one eye of the subjects at least thirty minutes before the start of the imaging session. These eyedrops caused dilation of the pupil (mydriasis) and inhibition of accommodation (cycloplegia), which are necessary to achieve sufficient resolution and image quality. Subjects sat in front of the AOSLO instrument and placed their chin on a chinrest or used a dental impression bar. Temple pads were then adjusted to immobilize the head and ensure the subjects’ stability in the system. A manual 3-axis translation stage was used to adjust the head position to keep the eye pupil centered on the illumination beam.

During the imaging session, subjects fixated on a 4.7 x 4.7 arcmin dark square presented within the 60 x 60 arcmin imaging field of view. This fixation marker was generated by modulating the AOM for the imaging channel, resulting in a small blinking dark square appearing on the red background, which was either 680 nm or 840 nm light. The fixation marker was flashed at 3 Hz with a 50% duty cycle. Subjects were instructed to fixate on the dark square, which was placed near the center of the imaging field of view. The power levels used during the imaging session were well below the retinal exposure limits set by the ANSI standard in Z136.1-2014 [33] (i.e., 32 μW for 680 (imaging), 74 μW for 840 nm (imaging) and 75 μW for 940 nm (wavefront sensing)). Imaging sessions were between 30 and 60 minutes in length. Multiple 10-second recordings were collected for each subject.

### 4.2 Data analysis methods

All data analysis was conducted offline after the imaging session was completed. Each 10-second recording was first manually graded by the experimenter to assess various factors contributing to imaging quality (i.e., average pixel intensity, sharpness of the retinal image, and fixational stability) and select the best recordings. The selected videos were stabilized using a strip-based image registration algorithm to compensate for retinal image motion during fixation [31,34,35]. The implementation of this registration technique was done using the software REMMIDE from the Rossi Lab [36]. REMMIDE uses a 3-step procedure to construct a low-distortion synthetic reference frame, eye motion traces, and a high-SNR stabilized retinal image. The stabilized retinal image was then contrast enhanced using adaptive histogram equalization in MATLAB. The retinal images with the best image quality—based on assessing the sharpness of the smallest foveal cones [36]—were then cropped with a 1-degree square window centered on the highest-density region of the fovea, as shown in column 1 of Fig. 8.

**Fig. 8:**
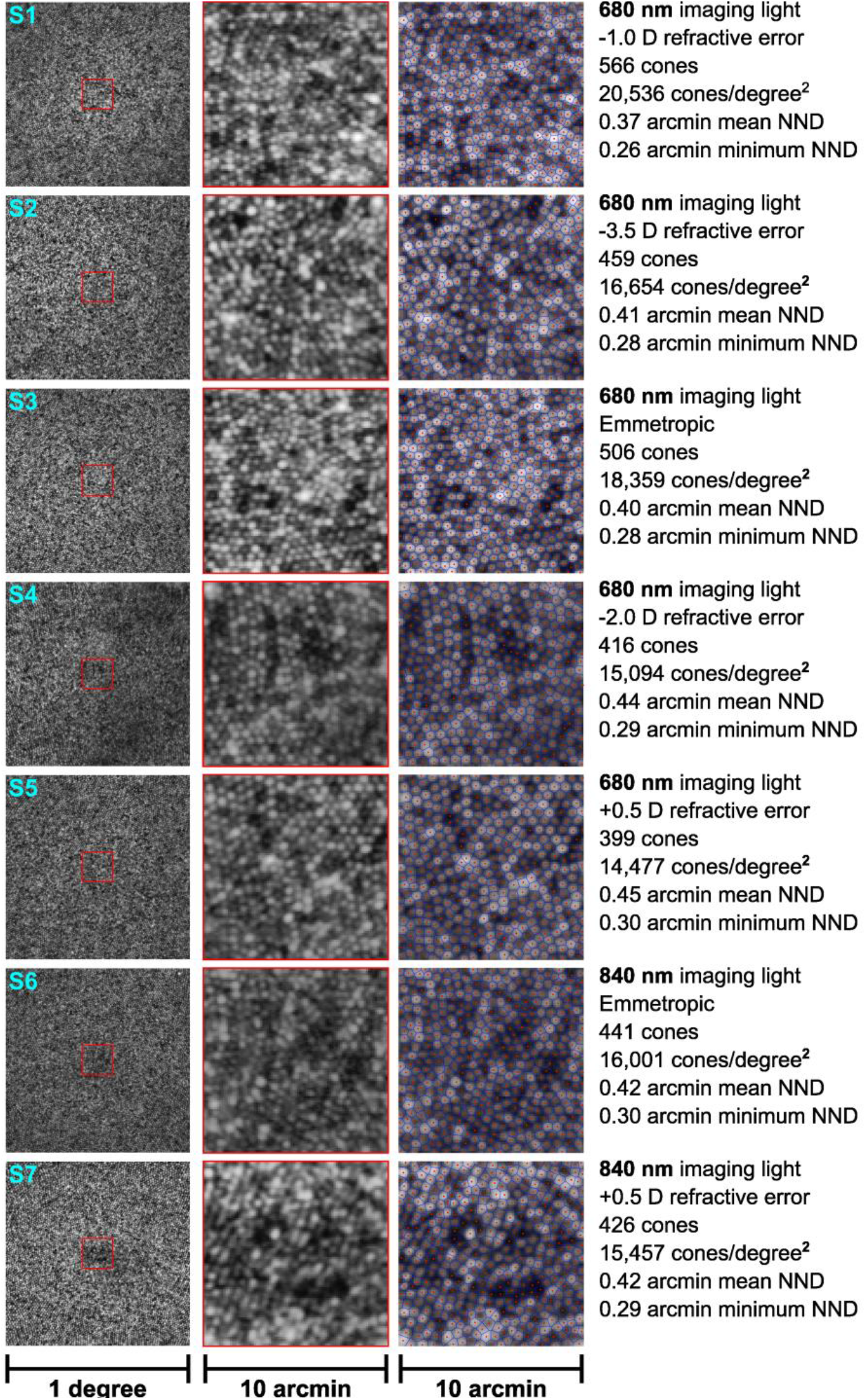
Images of the human photoreceptor mosaic at the center of the fovea for seven subjects collected with the aligned and optimized AOSLO using either 680 nm or 840 nm illumination. Column 1 shows a 1-degree square image centered on the highest cone density region. Column 2 shows a magnified view of the highest cone density region. Column 3 shows the same magnified view with cone centers marked (red dots) and a Voronoi diagram overlay (blue lines) denoting the cone borders. The fourth column summarizes the imaging conditions and the cone statistics. NDD refers to the cone nearest neighbor distance.

A 10-arcmin square window at the center of each retinal image was then selected for cone tagging, as shown in column 2 of Fig. 8. Cone centers were manually marked by a single observer. In cases where cones appeared dark, markers were still added at these locations. The decision to label these poorly reflective cones was based on the evidence that cone reflectivity can change between imaging sessions and low-reflectivity cones have normal detection thresholds [37]. When marking these dark cones, the regularity of the mosaic was prioritized based on histological measurements of the cone mosaic [38], as is common practice for cone tagging of AOSLO images [5,7]. Column 3 of Fig. 8 shows the results of the cone tagging, with red markers denoting the cone centers and blue lines showing the approximate cell boundaries using a Voronoi diagram overlay.

Quantitative analysis of the cone mosaic was conducted in angular units, with cone density reported in cones per square degree and nearest-neighbor distance (NDD) reported in arcmin. To compute the cone density, the number of cones in the 10-arcmin square window was divided by the window area. The cone density, along with the mean and minimum NND, is reported in Fig. 8 for each of the seven subjects. Axial length measurements with the ZEISS IOLMaster 700 were conducted for five of the seven subjects to compute a retinal magnification factor (RMF) for converting between angular units and length units on the retina following the methods of Bennett *et al*. [39]. The RMF values for the five subjects ranged from 0.271 to 0.293 millimeters per degree. Analyzing the cone NND data allowed for the angular and lateral resolution of the AOSLO to be quantified during human retinal imaging.

### 4.3 Results

Human retinal images captured with the AOSLO demonstrate cellular resolution in the center of the fovea. The mean cone NND ranged from 0.37 arcmin to 0.45 arcmin across the seven subjects (1.8 - 2.1 μm), with an average NND of 0.42 arcmin. Subject S1 had the highest cone density of 20,536 cones/degree^2^ and the smallest NND of 0.37 arcmin (1.8 μm). The AOSLO successfully resolved the smallest foveal cones for all seven subjects. Subjects S1-S5 were imaged with 680 nm light, while subjects S6 and S7 were imaged with 840 nm light. The reduced resolution of the 840 nm imaging channel did not preclude the smallest foveal cones from being resolved for these two subjects, but for subjects with higher cone densities (i.e., S1 and S3), the higher resolution of the 680 nm channel was necessary to resolve the smallest cones. The diffraction-limited resolution of the AOSLO is 0.40 arcmin (1.9 μm) for 680 nm light and 0.49 arcmin (2.4 μm) for 840 nm light. These imaging results further validate the resolution of the system, showing that the diffraction-limited resolution that was obtained with a resolution target continues to hold for the far more challenging recording conditions provided by the human retina.

## 5. Conclusion

Implementing an AOSLO for resolving the smallest cones in the human retina requires a well-aligned optical system. This study describes rigorous alignment, calibration, and validation procedures that enable development and maintenance of a well-tuned device. All procedures were described in thorough detail to enable step-by-step replication by laboratories across different disciplines.

Starting with a CAD model for the system and a laser-cut stencil, the coordinates for each optical component and corresponding mechanical mount were transferred to an optical table, enabling rapid assembly and coarse alignment of the optical system. Following this coarse alignment, an active alignment strategy utilizing a portable Shack-Hartmann wavefront sensor was employed to correct residual defocus in each of the afocal relay telescopes. By measuring the wavefront curvature at intermediate pupil planes and using principles and equations from first-order optics, the required axial shifts necessary to correct the residual defocus were determined and implemented.

Following this two-stage alignment procedure, various validation and calibration experiments were conducted to demonstrate the performance of the completed system. Wavefront measurements at the eye pupil plane demonstrated diffraction-limited performance for each of the four spectral channels. The wavefront sensor was then used to verify that the correct amount of defocus had been added to each channel to compensate for the longitudinal chromatic aberration of the human eye. Images of a distortion grid target were used to calibrate the scanning field of view and digitally correct for the sinusoidal distortion from the resonant scanner. Images of a resolution test target enabled the system resolution to be quantified for each imaging channel. Finally, human retinal imaging was conducted to demonstrate that the smallest foveal cones can be resolved in healthy subjects with normal vision.

The procedures described above provide a systematic approach for implementing high resolution AOSLO systems. This approach is especially valuable for resolving cones in the central fovea. By measuring and quantifying performance metrics—such as diopters of residual defocus or RMS wavefront error—during the alignment of the AOSLO, each optical subsystem can be individually optimized, resulting in a well-aligned and high-performing completed system. This work seeks to enable research groups to build AOSLOs and incorporate high-resolution human retinal imaging into their research efforts by describing a set of procedures covering many aspects of the AOSLO system implementation and validation. These methods were developed for applications where cellular resolution at the center of the fovea is necessary, but other AOSLO systems with less stringent resolution requirements—e.g., imaging over a larger field of view or outside of the fovea—will also benefit from the rigorous and systematic alignment and validation procedures presented.

The AOSLO described in this paper is currently being used in several human retinal imaging and psychophysics studies investigating the relationships between anatomical features of the cone photoreceptor mosaic and oculomotor behavior. Given the high quality of the images captured with this system, the cone locations can be identified using semi-automated cone detection procedures [40–42]. Cone density analysis is then conducted to map the cone density across the fovea and determine the location of the peak cone density relative to other foveal specializations. In psychophysics experiments, the location of the stimulus on the retina can be unambiguously determined through analysis of the recorded videos. Retinal image motion introduced by fixational eye movements can be measured through analysis of the recorded videos [36,43], providing another method for studying fixational eye movements at high resolution alongside DPI eye trackers and eye coils [44]. In addition, gaze-contingent stimulus presentation is also possible using the online stabilization capabilities of the imaging software [30,34,35]. These features of high-resolution AOSLO provide the capabilities for studying the acuity limits of human vision [45], the impact of fixational eye movements on visual acuity [46], and have the potential to shed light on how the interplay between anatomy, optics, and oculomotor behavior shapes visual acuity.

## Acknowledgements

The authors would like to acknowledge Ashley M. Clark, Sanjana Kapisthalam, Samantha K. Jenks, and Edith Hartmann for their assistance during the human retinal imaging sessions. We thank Sanam Mozaffari and Francesco LaRocca at UC Berkeley for making Zemax and SolidWorks designs of the AOSLO system available. William S. Tuten at UC Berkeley provided some custom hardware necessary to assemble the AOSLO. The Center for Visual Science at the University of Rochester (partial funding through NIH grant EY001319) provided resources and expertise valuable for completing this research, especially Martin Gira and Daniel Guarino for fabricating custom mechanical components and Karteek Kunala for advice on optical alignment techniques. Mike Pomerantz helped with the laser-cut stencil and the assembly of the custom Shack-Hartmann wavefront sensor. We thank Synopsys, Inc. for the student license of CODE V and Onshape for the student license for their CAD modeling software.

## Data availability

Data files underlying the results in this paper are available upon reasonable request.

